# A guide for membrane potential measurements in Gram-negative bacteria using voltage-sensitive dyes

**DOI:** 10.1101/2022.04.30.490130

**Authors:** Jessica A. Buttress, Manuel Halte, J. Derk te Winkel, Marc Erhardt, Philipp F. Popp, Henrik Strahl

## Abstract

Transmembrane potential is one of the main bioenergetic parameters of bacterial cells, and is directly involved in energising key cellular processes such as transport, ATP synthesis, and motility. The most common approach to measure membrane potential levels is through use of voltage-sensitive fluorescent dyes. Such dyes either accumulate or are excluded from the cell in a voltage-dependent manner, which can be followed by means of fluorescence microscopy, flow cytometry, or fluorometry. Since the cell’s ability to maintain transmembrane potential relies upon low membrane ion conductivity, voltage-sensitive dyes are also highly sensitive reporters for the activity of membrane-targeting antibacterials. However, the presence of an additional membrane layer in Gram-negative (diderm) bacteria significantly complicates their use. In this manuscript, we provide guidance on how membrane potential and its changes can be reliably monitored in Gram-negatives using the voltage-sensitive dye DiSC_3_(5). We also discuss the confounding effects caused by the presence of the outer membrane, or by measurements performed in buffers rather than growth medium. We hope that the discussed methods and protocols provide an easily accessible basis for the use of voltage-sensitive dyes in Gram-negative organisms, and raise awareness of potential experimental pitfalls associated with their use.

## INTRODUCTION

Due to their misuse and overuse in both clinical and agricultural settings, antibiotic resistance is one of the biggest threats to global health today. This crisis is exacerbated by a deficit in antibiotic innovation, as demonstrated by the linear decline in discovery of new antibacterial molecules over the past 30 years [1]. There is, therefore, an urgent need for compounds with novel targets and modes of action. One emerging strategy is targeting bacterial membranes [2].

The bacterial cytoplasmic membrane is an essential macromolecular structure that harbours critical cellular processes such as nutrient and waste transport, respiration and ATP synthesis, protein secretion, motility, cell division, and cell wall synthesis [3]. Hence, maintaining the cytoplasmic membrane in an intact, biologically functional, and selectively permeable state is critical for the viability of bacterial cells. One of the essential features of the bacterial cytoplasmic membrane is its ability to maintain an electrical potential (transmembrane potential) which, alongside ATP, is a key cellular energy reserve used to drive important energy-demanding processes such as ion homeostasis, nutrient, protein and lipid transport, ATP synthesis, and motility.

The suitability of bacterial membranes as a vulnerable drug-target is perhaps best demonstrated by eukaryotic host defence peptides, which act by disrupting bacterial membranes and play a critical role in our innate immune system [4–6]. Despite a long history of co-evolution, bacteria have failed to evolve resistance mechanisms that fully protect themselves against membrane-targeting host defence peptides. More recently, the clinical efficacy of membrane-active antimicrobials has been demonstrated by the success of polymyxin B, daptomycin and colistin as last resort antibiotics used to treat life-threatening infections caused by multi-drug resistant pathogens [7–10]. Whilst the Gram-negative outer membrane and the associated lipopolysaccharide (LPS) layer are formidable barriers against many agents that target the cytoplasmic membrane, polymyxin B and colistin show selective activity against Gram-negative bacteria. Additionally, host defence peptides of the Cathelicidin type are capable of disturbing the Gram-negative outer membrane as part of their antibacterial mode of action [4]. Hence, targeting the cytoplasmic membrane is a reasonable drug-development strategy, even against more challenging Gram-negative pathogens.

Membrane-targeting antibiotics commonly perturb membrane integrity by increasing permeability to ions or larger molecules, or by inducing more subtle changes such as forming or disturbing lipid domains, altering membrane fluidity, or delocalising membrane-associated proteins [11–15]. Large membrane-impermeable fluorescent dyes such as Propidium Iodide and Sytox Green are most frequently used to investigate antibiotic-induced changes in membrane permeability *in vivo.* These probes are DNA-intercalating and stain the nucleoid when large pores are formed in the cytoplasmic membrane, or when cell lysis is induced [4, 16, 17]. However, these assays are unable to detect smaller-sized channels, increased ion permeability, or inhibition of respiration; all of which can still be lethal to the cell through dissipation of the transmembrane potential and associated cellular consequences [11, 18–21]. Membrane depolarisation can be followed more directly using a number of voltage-sensitive fluorescent probes including 3,3’-Dipropylthiadicarbocyanine iodide DiSC_3_(5). Due to its hydrophobic and cationic nature, DiSC_3_(5) can penetrate the lipid bilayer and accumulate to high levels in polarized cells; a process which is associated with self-quenching of fluorescence. Upon membrane depolarisation, the dye is rapidly released from the cells resulting in a dequenching which can be followed fluorometrically, microscopically, and using flow cytometry [22, 23]. Whilst DiSC_3_(5), and a closely related voltage-sensitive dye 3,3’-Diethyloxacarbocyanine Iodide (DiOC_2_(3)), are frequently used in antibiotic mechanism-of-action studies in Gram-negative bacteria [24–26], the used protocols are relatively inconsistent and frequently include incubations in buffers of various compositions, presence of chelating agents such as EDTA, or in strains with hyperpermeable outer membranes. Accordingly, robust protocols for using these voltage-sensitive dyes to measure perturbations of the transmembrane potential in Gram-negative bacteria are missing.

Reliable and reproducible assays to follow changes in bacterial membrane potential are not only important in the context of antibiotic research. The use of DiSC_3_(5) as proxy for the polarisation state of cells could be applied to study distinct cellular processes such as the energetic burden of flagellar formation and rotation. The assembly of the bacterial flagellum, as well as the flagellar-mediated swimming motility both rely on the proton motive force [27, 28]. Recently, proton leakage via the stator units of the flagellar motor has been associated with a reduced growth rate in *Salmonella* [29]. Thus, voltage-sensitive dyes, such as DiSC_3_(5), could be used to monitor the effects of such energy-consuming processes on the polarisation state of individual cells and their current physiological condition.

In this manuscript, we will discuss and provide guidance on fluorescence-based techniques to measure membrane potential in the Gram-negative model organisms *Escherichia coli* and *Salmonella enterica.* We hope to raise awareness regarding the various confounding parameters and factors that can have a significant effect on the voltage-dependent behaviour of dyes in the context of Gram-negative bacteria. The discussed methods and protocols should provide a useful starting point for colleagues interested in identifying and characterising the antibacterial mode-of-action of membrane-targeting compounds against Gram-negative bacteria, or for analysing Gram-negative membrane potential levels in a more physiological context.

## METHODS

### Strains, media and growth conditions

Strains and genotypes are listed in Table 1. *E. coli* was grown in Lysogeny Broth (Miller) [10 g/l tryptone, 5 g/l yeast extract, 10 g/l NaCl] and *S. enterica* in Lysogeny Broth (Lennox) [10 g/l tryptone, 5 g/l yeast extract, 5 g/l NaCl]. For experiments performed in buffer, cells were collected by 3 min centrifugation at 6000×g, washed, and resuspended to an OD_600_ of 0.3 in phosphate-buffered saline (PBS) [8 g/l NaCl, 0.2 g/l KCl, 1.15 g/l Na_2_HPO_4_, 0.2 g/l KH_2_PO_4_, pH 7.3). If indicated, PBS was supplemented with 0.2% glucose and 1 mM CaCl_2_.

**Table 1:** Strains used in this study

| Strain | Genotype | Reference |
| --- | --- | --- |
| *E. coli* MG1655 | $\lambda^+$ F<sup>-</sup> *ilvG*- *rfb*-50 *rph*-1 | [48] |
| *S. enterica* TH437 | *Salmonella enterica* serovar Typhimurium strain LT2 | [49] |

### Minimal inhibitory concentration (MIC) determination

*E. coli* overnight cultures were diluted 1:100 in appropriate growth medium and grown to mid-logarithmic phase. Cells were then diluted to give a final concentration of 5 × 10^5^ cells per well in a pre-warmed 96-well microtiter plate. This plate was prepared with an initial high concentration of the desired compound followed by a serial 2-fold dilution. After addition of the cells, the plate was incubated at 37°C for 16 hours with shaking at 700 rpm. MIC was defined as the lowest compound concentration able to inhibit visible bacteria growth.

### Fluorescence microscopy

*E. coli* overnight cultures were diluted 1:100 in LB and incubated at 37°C upon shaking until an OD_600_ of 0.3. 100-200 μl culture aliquots were transferred to 2 ml Eppendorf tubes with perforated lids followed by addition of 7 μM (10 μg/ml) polymyxin B (Sigma-Aldrich) or 31 μM (30 μg/ml) polymyxin B nonapeptide (Sigma-Aldrich) and incubation at 37°C upon shaking using a thermomixer. If applicable for the specific experiment, 200 nM of the membrane permeability indicator Sytox Green solved in H_2_O (ThermoFisher) was added alongside the antimicrobial compound, whilst 1 μM of the membrane potential-sensitive dye DiSC_3_(5) (Sigma-Aldrich) was added 5 min prior to imaging. The DMSO concentration of the final cell suspension was kept at 1-2%, which is critical for good DiSC_3_(5) solubility and staining while not affecting growth itself. Samples were immobilised on Teflon-coated multi-spot microscope slides (ThermoFisher) covered with a thin layer of H_2_O/1.2% agarose and imaged immediately. In case of data shown in Fig. 1, agarose was additionally supplemented with 10% LB. For further details about this type of slide preparation, see te Winkel *et al.* [23]. Microscopy was performed using a Nikon Eclipse Ti equipped with Nikon Plan Apo 100×/1.40 Oil Ph3 objective, CoolLED pE-4000 light source, Photometrics BSI sCMOS camera, and Chroma 49002 (EX470/40, DM495lpxr, EM525/50) and Semrock Cy5-4040C (EX 628/40, DM660lp, EM 692/40) filter sets. Images were acquired with Metamorph 7.7 (MolecularDevices) and analysed with Fiji [30].

**Figure 1:**
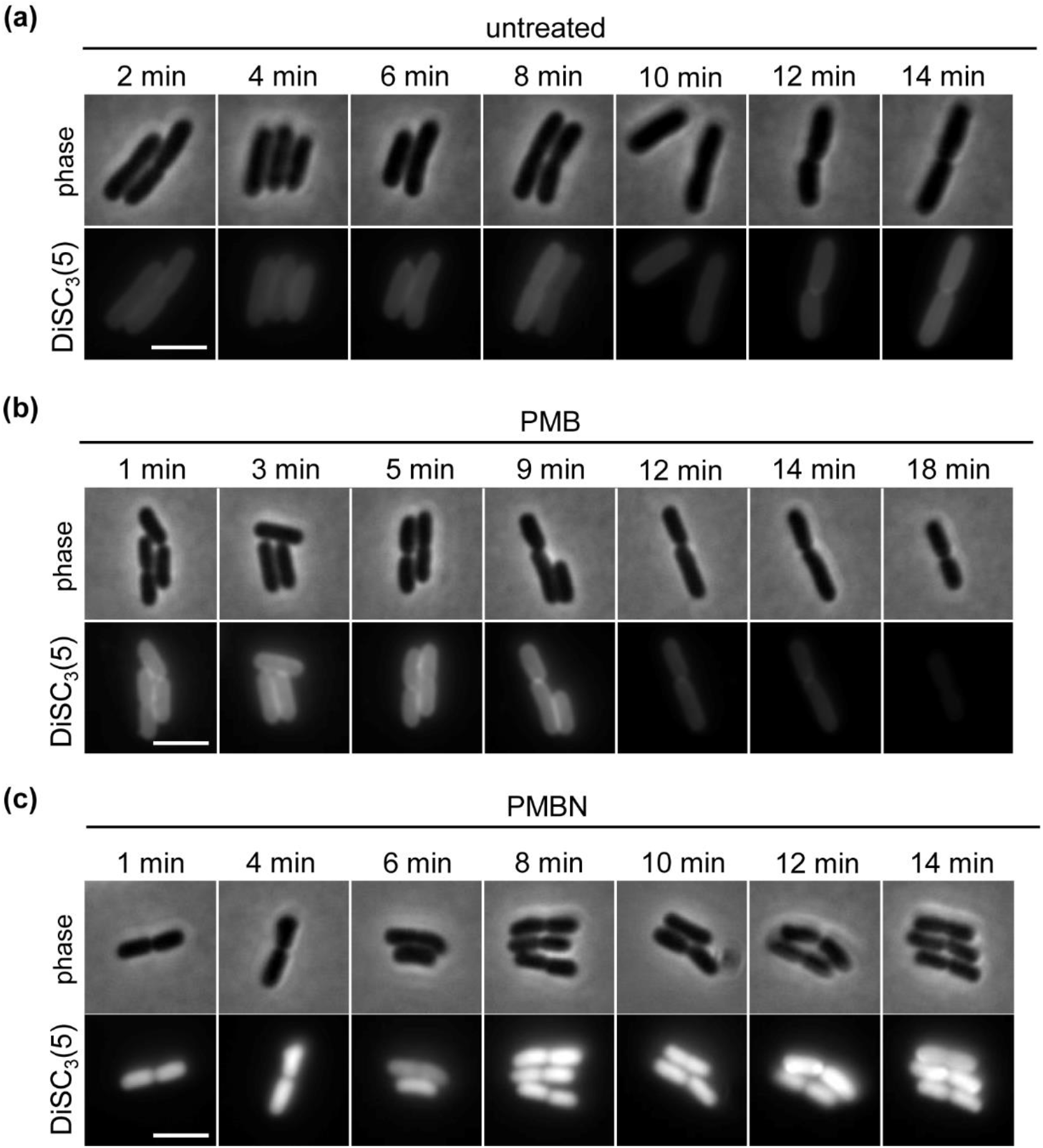
*E. coli* DiSC_3_(5) staining is influenced by both outer membrane permeabilisation and (inner) membrane depolarisation. Phase contrast and fluorescence microscopy of *E. coli* stained with DiSC_3_(5) in the (a) absence and presence of (b) outer membrane permeabilising and inner membrane depolarising PMB (7 μM) or (c) outer membrane permeabilising PMNBN (30 μM) at different time points of incubation. Note that for this experiment the dye and antibiotics were added directly to the agarose pad supplemented with 10% LB. The time points indicate incubation after transfer of cells to the agarose pad. Scale bar: 3 μm. Strain used: *E. coli* MG1655 (wild type).

### Time-lapse microscopy

*E. coli* wild type cells were grown overnight at 30°C in M9 medium supplemented with 0.4% glucose, 0.2% casamino acids and 1mM thiamine, followed by a 1:10 dilution in M9 medium with diluted nutrients (0.02% glucose and 0.01% casamino acids) and incubation at 30°C. The time-lapse slide was prepared as described earlier [23, 31] with following modifications. For the preparation of the slide, a 2× stock of carbon source-diluted M9 medium (0.02% glucose, 0.01% casamino acids) was preheated to 60°C and diluted 2-fold with 3% low-melting agarose kept at 60°C. DiSC_3_(5) solved in DMSO was added to the agarose to a final concentration of 2.5 μM and a DMSO concentration of 1%, followed by pouring the time-lapse slide. The cells growing in nutrient arm M9 liquid medium were diluted to an OD_600_ of 0.035, followed by transfer to the time lapse slide. Time-lapse microscopy was carried out at 30°C using an Applied Precision DeltaVision RT automated microscope equipped with a Zeiss Plan Neofluar 63×/1.40 Oil Ph3 objective and Photometrics CoolSnap HQ camera, and a Cy5 filter set (EX632/22, DM645-700, EM 679/34). Phase contrast and fluorescence images were taken every 10 minutes over the duration of the microcolony growth (19h).

### Image analysis

Quantification of DiSC_3_(5)-fluorescence for individual cells was performed in a semi-automated manner using Fiji [30]. To eliminate bias, the individual cells were identified and converted to regions of interest (ROIs) by thresholding of corresponding phase contrast images. If individual cells adhered to each other in a clearly identifiable manner, they were manually separated prior to automated cell detection. Larger cell aggregates were omitted from the analysis. The fluorescence intensity values for individual cells across the population were obtained from background-subtracted fluorescence images by using the phase contrast imaging-derived ROIs. To further eliminate bias, only multiples of full fields of view were analysed.

### Fluorometric determination of membrane potential levels

#### E. coli

Cultures were grown to logarithmic growth phase and, if needed, diluted to an OD_600_ of 0.5 in growth medium supplemented with 0.5 mg/ml fatty acid free BSA (Sigma-Aldrich). Addition of BSA is critical to supress DiSC_3_(5) binding to microtiter plate plastic surface. Immediately prior to measurement, the cells were transferred to black polystyrene 96-well plates (Porvair Sciences) and the autofluorescence of *E. coli* was measured for up to 5 min. DiSC_3_(5) dissolved in DMSO was then added to a final concentration of 0.5 μM (1% DMSO) and the fluorescence quenching was monitored until a stable baseline was obtained, followed by addition of 7 μM (10 μg/ml) polymyxin B (Sigma-Aldrich) or 31 μM (30 μg/ml) polymyxin B nonapeptide (Sigma-Aldrich). Fluorometric measurements were taken every minute, with vigorous shaking in between readouts, using a BMG Clariostar multimode plate reader upon 610 nm (± 10) excitation, and 660 nm (± 10) emission settings. All media, plates and instruments were warmed to 37°C prior to use. To investigate whether compounds of interest interfered with DiSC_3_(5) fluorescence at used concentrations, this assay was repeated, in the absence of cells, in PBS supplemented with BSA.

#### S. enterica

For determination of membrane potential of stationary phase cells, the OD_600_ of overnight cultures was determined and subsequently 2 ml were harvested by centrifugation for 3 min at 21,000×g to obtain cell free spent medium. To prevent re-energisation though resuspension in fresh medium, the cells were diluted to an OD_600_ of 0.2 in spent medium, mixed with 0.5 mg/ml fatty acid free BSA (Sigma-Aldrich) and incubated in a thermomixer for 15 min at 850 rpm and 37°C with open lids. To pre-treat samples, polymyxin B nonapeptide (Merck) solved in H_2_O was added to a final concentration of 31 μM (30 μg/ml). After that, cell suspension was transferred into black 96-well polystyrene plates (Greiner Bio-One) and the autofluorescence of *S. enterica* was recorded at 610 nm (± 9) excitation and 660 nm (± 20) emission every minute for 3 min using a Tecan Infinite M200 plate reader with continuous shaking between the measurements. Subsequently, DiSC_3_(5) (Eurogentech) was added to a final concentration of 1 μM while maintaining 1% DMSO, and fluorescence measurement was continued for another 17 min in one minute intervals. For pre-treated samples, polymyxin B (Merck) was added to a final concentration of 14 μM (20 μg/ml), whilst water was added for untreated controls. For non-pre-treated samples, either polymyxin B nonapeptide or polymyxin B were added to final concentrations of 31 μM (30 μg/ml) or 14 μM (20 μg/ml), respectively. Membrane potential was followed for another 30 min in the plate reader with readings every minute. For measuring membrane potential of exponential growth phase growing cells, a fresh sub-culture was inoculated 1:100 from an overnight culture and incubated until an OD_600_ between 0.3 and 0.7 was reached. Subsequently, cells were diluted in fresh medium to an OD_600_ of 0.2 and the protocol was performed as described above.

### Statistical analyses

The data presented here are representatives of at least two independent experiments. The minimal inhibitory concentration (MIC) and fluorometric assays were carried out as technical triplicates. The statistical significance was calculated as one-way, unpaired ANOVA with Tukey’s post hoc test. Significance was depicted as **** for p < 0.0001, *** for p < 0.001, ** for p < 0.01, * for p < 0.05, ns for p > 0.05.

## RESULTS AND DISCUSSION

### Inhibitory effects of used dyes and compounds

A valid concern when using voltage-sensitive and other cell dyes is their potential to alter the cellular properties they are intended to monitor. Indeed, the amyloid [32], RNA [33, 34], and voltage-sensitive dye [35, 36] thioflavin T (ThT) was recently shown to dissipate the membrane potential itself if used in too high concentrations [37]. To test the growth inhibitory potential of DiSC3(5), and of the antimicrobial peptides used throughout this study, standard minimum inhibitory concentration (MIC) assays were carried out with *E. coli* MG1655 and *S. enterica* LT2, used here as wild type strains. As shown in table 2, Polymyxin B (PMB) displayed strong antibacterial activity against both species. This, along with its well-documented ability to permeabilise both the outer and the cytoplasmic membranes [7, 38], confirmed its validity as a suitable positive control for this study. Crucially, PMB does not exhibit dye-interactions with DiSC_3_(5) (Fig. S1); a problematic phenomenon that frequently occurs between hydrophobic dyes and antimicrobials; which we recommend to always test and rule out [23]. As expected, a nonapeptide derivative of Polymyxin B (PMBN), which retains the ability to disrupt the Gram-negative outer membrane but is unable to form the depolarising cytoplasmic membrane pores [38], did not inhibit growth at any concentration tested (Table 2).

**Table 2:**
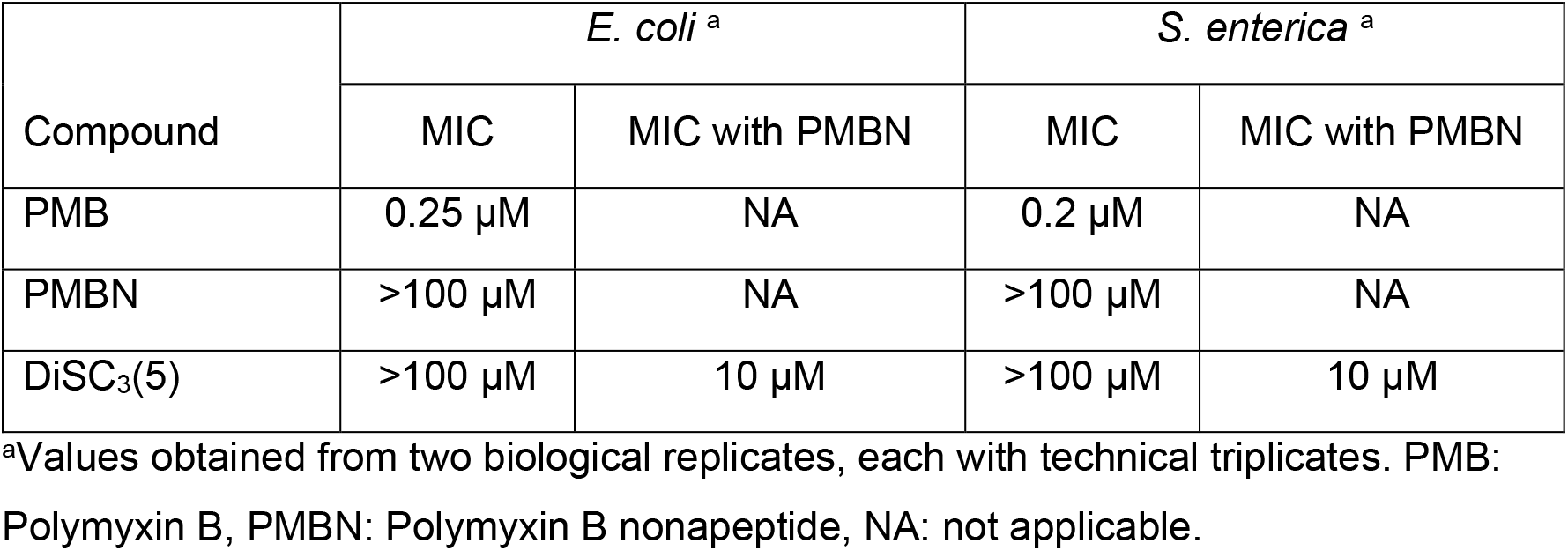
Minimum inhibitory concentrations (MICs) of tested compounds and dyes for *E. coli* and *S. enterica* in the presence and absence of the outer membrane permeabilising agent Polymyxin B nonapeptide.

Previously, we have reported that DiSC_3_(5) is growth-inhibitory in the Gram-positive model organism *Bacillus subtilis* through a mechanism that is currently unknown but that does not involve membrane depolarisation [23]. However, it appeared that this inhibitory effect is a peculiarity of *B. subtilis* and does not occur in the close relative *Staphylococcus aureus,* or the Gram-negative species used in the present study (Table 2). When combined with the outer membrane permeabilising compound PMBN, an increased sensitivity of DiSC_3_(5) was observed with a MIC of 10 μM (Table 2), which is higher than the concentrations applied in subsequent experiments. We verified this by applying 1 μM of DiSC_3_(5) to *E. coli* cells in logarithmic phase and observed no effect on growth even in the presence of PMBN (Fig. S2).

### DiSC_3_(5) staining is influenced by both (inner) membrane potential and outer membrane permeabilisation

As shown previously, single-cell DiSC_3_(5) fluorescence microscopy provides a simple and rapid method to measure membrane potential in the Gram-positive model organism *B. subtilis* [23]. To test whether this approach is also feasible in Gram-negative bacteria, we analysed the staining of wild type *E. coli* cells in the absence and presence of polymyxins (PMB and PMBN). For this aim, we included DiSC_3_(5) and the respective peptides in agarose pads, additionally supplemented with 10% LB, followed by addition of *E. coli* cells and rapid microscopy. This was to observe the coarse kinetics of membrane depolarisation upon the imaging process. In untreated cells, DiSC_3_(5) fluorescence signals remained stable over 14 min (Fig. 1). Surprisingly, an immediate increase in DiSC_3_(5) fluorescence intensity for approx. 10 min was observed upon incubation with PMB, which was then followed by a loss of DiSC_3_(5) signal indicating membrane depolarisation. This increase was even more apparent upon PMBN-treatment where the cells remained highly stained for the duration of the experiment (approx. 15 min). This demonstrates that whilst DiSC_3_(5) staining is sensitive to (inner) membrane potential levels and exhibits the expected loss of staining upon depolarisation, the staining is also strongly influenced by outer membrane permeabilisation. In conclusion, DiSC_3_(5) is indeed well-suited for detection of cytoplasmic membrane potential levels in wild type *E. coli* cells directly in growth medium. However, care should be taken when interpreting the results if used under conditions that compromise the integrity of the outer membrane, or when comparing strains that have outer membranes of different composition or structure.

### Membrane potential measurements in buffer are possible but problematic

Whilst DiSC_3_(5) has been previously used by our groups and others in the context of Gram-negative cells, the measurements are frequently carried out in cells washed with buffers of varying composition [25, 39, 40]. To investigate how washing and resuspending cells in buffer influences DiSC_3_(5) staining, we compared the DiSC_3_(5) signal levels between cells washed in phosphate-buffered saline (PBS) and cells stained directly in the growth medium. One important but frequently overlooked parameter when analysing membrane potential levels in buffer-suspended cells is the necessity to maintain a metabolisable carbon source. To test its effect on DiSC_3_(5) signal levels, we compared cells grown in LB supplemented with 0.2% glucose, and cells washed and resuspended in PBS with and without 0.2% glucose, followed by incubation for 15 min with shaking and staining. PMB-treatment for 15 min was used as a positive control for complete membrane depolarisation. The DiSC_3_(5) signal levels for cells washed in PBS with glucose were higher compared to cells in the growth medium and extremely heterogeneous at the single-cell level (Fig. 2a, b). In the absence of a carbon source, the heterogeneity was still present. However, as shown in both the microscopy images and quantification of single-cell fluorescence levels, the DiSC_3_(5) staining was greatly reduced upon carbon source withdrawal indicating strongly reduced membrane potential (Fig. 2a, b). Whilst the higher initial DiSC_3_(5) staining levels in PBS may be due to differences in solubility of the dye between growth medium and buffer, we hypothesised that washing the cells could also remove divalent cations that are critical for outer membrane stability [38, 41, 42]. Hence, PBS may slightly permeabilise the outer membrane, thus explaining the increased DiSC_3_(5) staining. To test this, we repeated the experiment washing and resuspending the cells in PBS additionally supplemented with 1 mM CaCl_2_. Again, signals were diminished in the absence of a carbon source (Fig. 2a, b). However, supplementation of PBS with both glucose and CaCl2 improved the consistency of signals and gave rise to DiSC_3_(5) fluorescence intensities more comparable to those measured in growth medium. This demonstrates that divalent cation removal, likely through destabilisation of the outer membrane, indeed affects DiSC_3_(5) staining.

**Figure 2:**
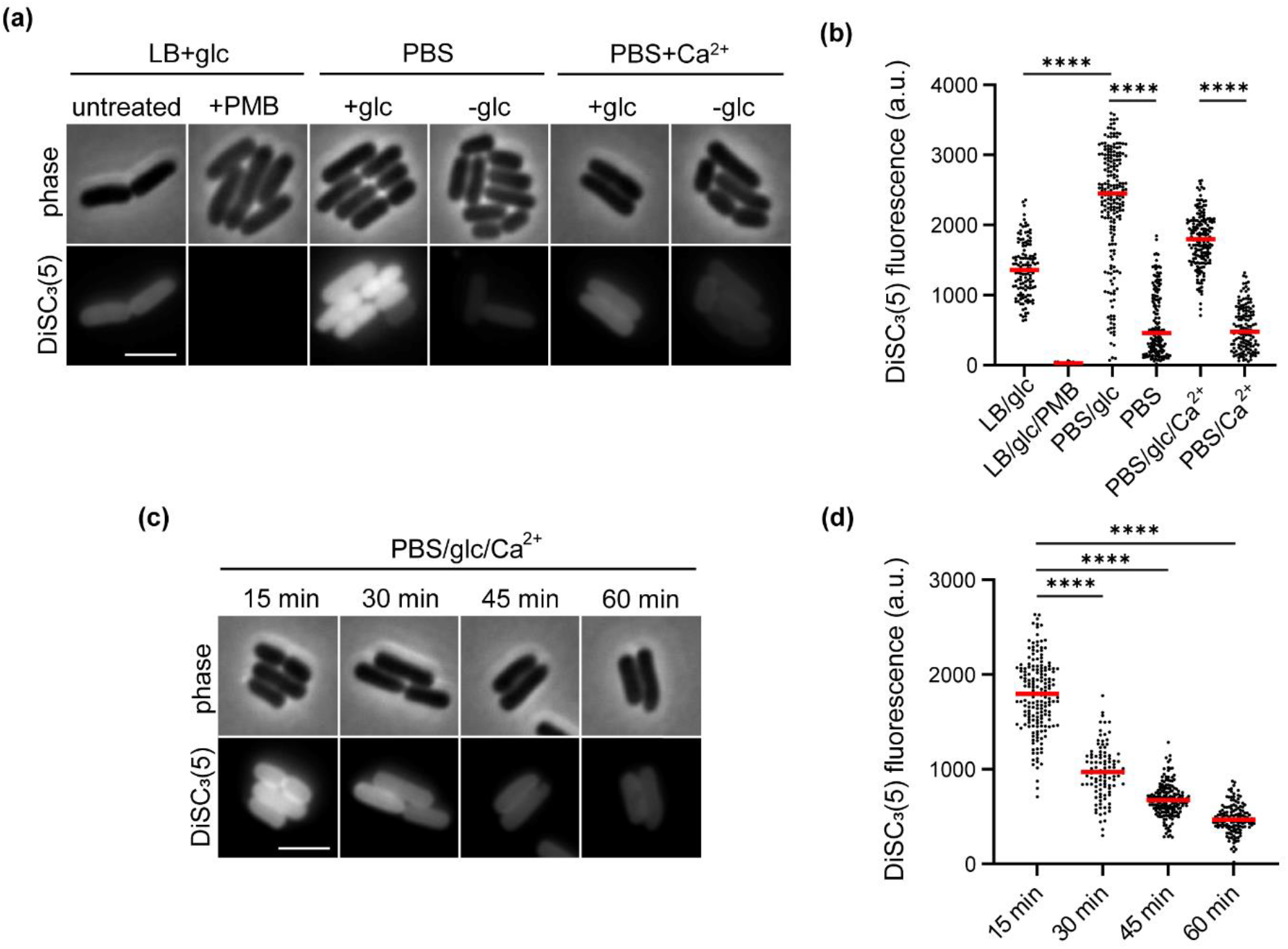
Divalent cations, carbon source, and rapid imaging are critical for measuring *E. coli* membrane potential in buffer. (a) Phase contrast and fluorescence microscopy images of DiSC_3_(5)-stained *E. coli* in LB/0.2% glucose, in PBS with and without glucose (0.2%), and in PBS/glucose (0.2%) with 1 mM CaCl2. As a positive control, the transmembrane potential was disrupted by the pore forming antibiotic PMB (7 μM). (b) Quantification of DiSC_3_(5)-fluorescence for individual cells from the imaging dataset shown in panel a (n=128-211 cells). Median fluorescence intensity is indicated with a red line, together with P values of a one-way, unpaired ANOVA with Tukey’s post hoc text. **** represents p < 0.0001. Scale bar: 3 μm. Strain used: *E. coli* MG1655 (wild type).

At last, we performed a DiSC_3_(5) fluorescence microscopy time course experiment with cells washed and resuspended in PBS to investigate how long they remain energised. As demonstrated by the loss of DiSC_3_(5) fluorescence, *E. coli* cells gradually loose membrane potential in PBS buffer even when supplemented with a carbon source and CaCl_2_ (Fig. 2c, d).

In conclusion, whilst DiSC_3_(5) can be used as a voltage-sensitive dye in buffers, assays carried out directly in the growth medium should be strongly favoured. If for experimental reasons measurements in buffer are essential, care must be taken to maintain both a carbon source to sustain central carbon metabolism and divalent cations to maintain outer membrane stability, and to carry out the assays rapidly after wash and resuspension into buffers.

### Compatibility of DiSC_3_(5) with time lapse microscopy and combination with other fluorophores

As DiSC_3_(5) is not growth inhibitory in *E. coli,* we hypothesised that it could be compatible with time-lapse experiments. To test this, we performed a time-lapse microscopy experiment with *E. coli* grown on DiSC_3_(5)-supplemented agarose pads using a method previously described in detail for other bacterial species [23, 31]. We chose to carry out this experiment in M9/glucose/casamino acids medium with both glucose and casamino acids concentrations reduced to 10% from normal. In this regime the growing microcolony exhausts the locally available carbon sources before exceeding the camera field of view. As shown in both Fig. 3 and Movie S1, DiSC_3_(5) fluorescence can indeed be monitored in a time-lapse microscopy setting for an extended duration of time. Crucially, the cessation of growth coincides with a strong reduction of DiSC_3_(5) fluorescence, consistent with membrane depolarisation triggered by carbon source exhaustion.

**Figure 3:**
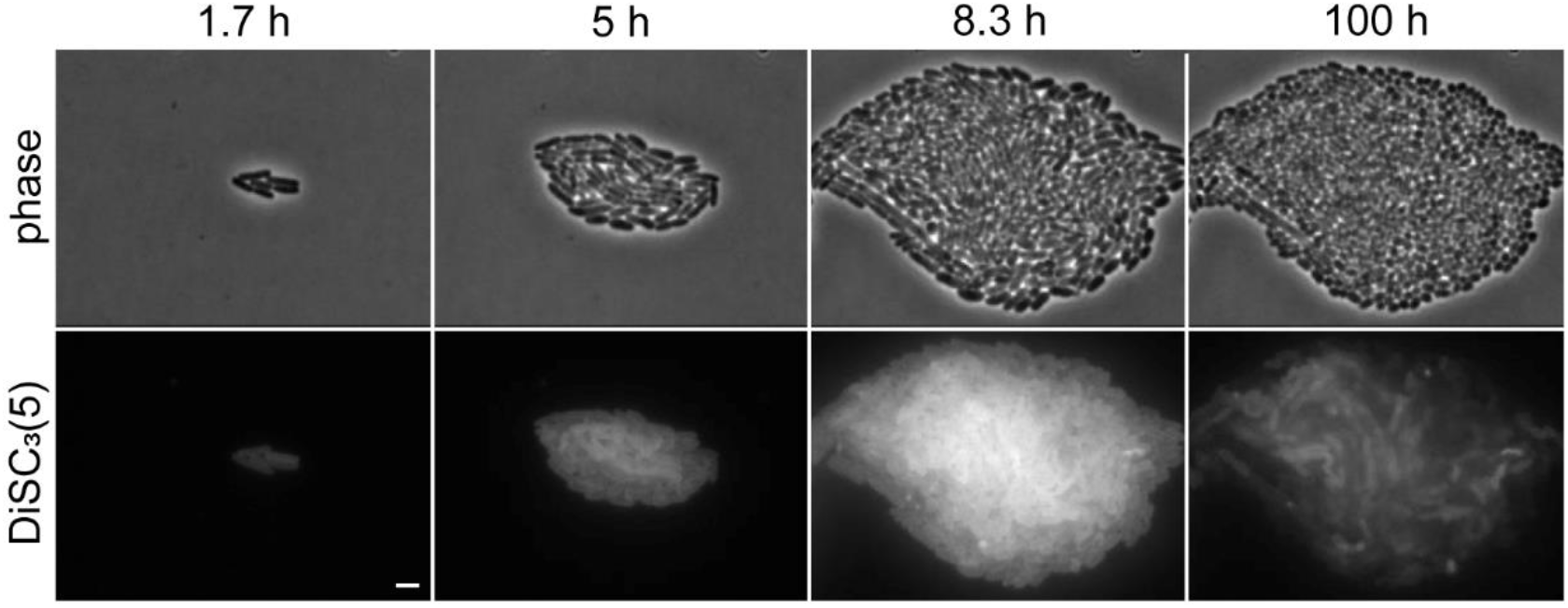
Time lapse microscopy of DiSC_3_(5)-stained *E. coli.* Selected phase contrast and fluorescence images of *E. coli* stained with DiSC_3_(5), and growing as a microcolony on M9-medium with limited carbon sources. Note the high DiSC_3_(5) staining in well-energised, actively growing cells and reduced staining upon entry into nutrient starvation-induced stationary growth phase indicating reduced membrane potential. The interior of the colony shows apparent higher staining dye to multiple cell layers rather than higher membrane potential. See Supplementary Movie 1 for the whole time-lapse series. Scale bar: 3 μm. Strain used: *E. coli* MG1655 (wild type).

Another useful property of DiSC_3_(5) is its far-red fluorescence emission spectrum (approx. 650-700 nm). While covered by commonly used Cy5-filters, it allows DiSC_3_(5) to be used in combination with even relatively weak green fluorophores such as GFP. This enables experiments that combine membrane potential readout with GFP-based protein localisation or expression reporter. One fluorophore combination that we have found to be very informative in the context of antibiotic research is co-staining with Sytox Green. Sytox Green is a membrane impermeable DNA intercalating dye that can stain the bacterial DNA but only when large pores are formed in the cytoplasmic membrane [16]. As shown in Fig. 4, *E. coli* can be simultaneously co-stained with both DiSC_3_(5) and Sytox Green. Upon treatment with the pore forming PMB, DiSC_3_(5) fluorescence is lost due to depolarisation whilst cells become strongly stained with Sytox Green indicating that the observed depolarisation is, as expected, caused by pore formation. This dual-dye technique thus enables the rapid fluorescence-based identification and differentiation between membrane depolarising and membrane pore-forming antimicrobial compounds or stresses *in vivo,* on a single-cell level.

**Figure 4:**
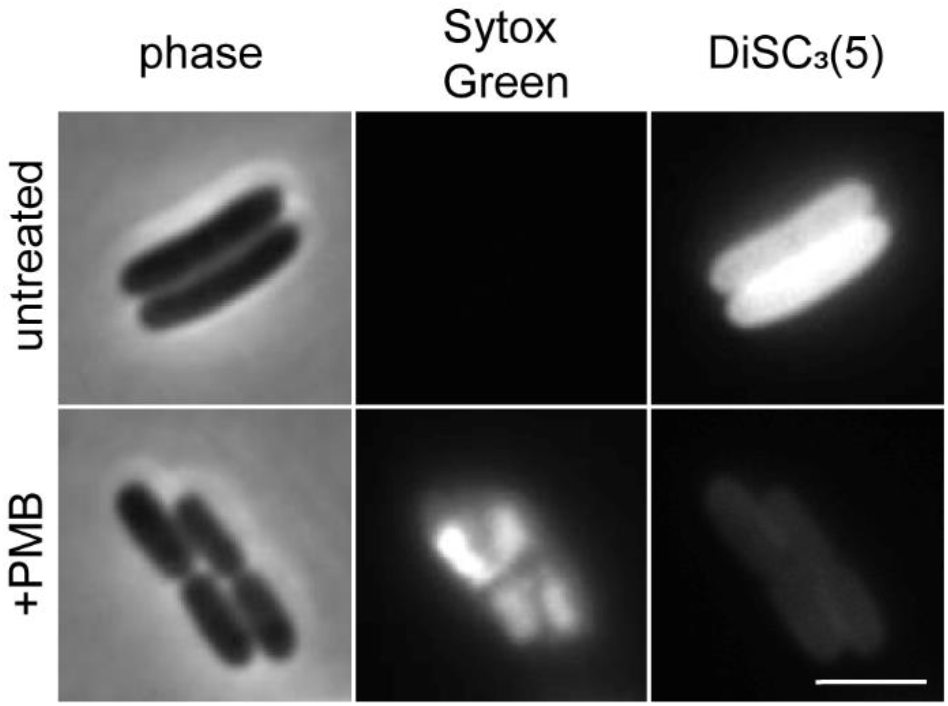
Simultaneous detection of membrane potential and pore formation in *E. coli.* Phase contrast and fluorescence microscopy of *E. coli* cells co-stained with Sytox Green and DiSC_3_(5) in the absence or presence of PMB (7 μM) for 15 min. Note the loss of membrane potential in PMB-treated cells that coincides with ability of Sytox Green to enter the cells indicating pore formation. Scale bar: 3 μm. Strain used: *E. coli* MG1655 (wild type)

### DiSC_3_(5)-based membrane potential measurements using fluorometry

The approaches detailed above allow *E. coli* membrane potential levels to be monitored at the single-cell level using fluorescence microscopy and flow cytometry. Whilst sacrificing single-cell resolution, DiSC_3_(5)-based membrane potential assays using fluorometry are perhaps more accessible, and provide better throughput and temporal resolution. This approach is based on self-quenching of DiSC_3_(5) fluorescence upon accumulation to high concentrations in polarised cells. When measured fluorometrically in a cell suspension, the accumulation of DiSC_3_(5) is observed as a gradual decline in fluorescence signal until a Nernstian equilibrium is achieved [43–45]. Upon loss of membrane potential, DiSC_3_(5) is released back into the medium, which leads to dequenching and an increase of overall measured fluorescence. Following DiSC_3_(5) fluorescence quenching behaviour, thus, enables live monitoring of mean membrane potential levels of a cell suspension. In the following, we will focus on such measurements for *S. enterica* cells exposed to the antimicrobials, PMB or PMBN.

The degree of DiSC_3_(5) fluorescence quenching is highly dependent on the used dye concentration and cell densities. Similar to the optimisation previously undertaken in Gram-positive bacteria [23], we first determined how the outer membrane barrier of Gram-negative bacteria affects the quenching of DiSC_3_(5) fluorescence upon accumulation in well energised cells. Upon addition of DiSC_3_(5) to exponentially grown but PMBN naïve cells, we observed only a slow and gradual quenching (Fig. 5a). When these cells were challenged with PMBN a further decrease in DiSC_3_(5) fluorescence was observed, indicating that outer membrane permeabilisation induced by PMBN stimulated further DiSC_3_(5) uptake, which is consistent with our microscopic observations in *E. coli* (Fig. 1). This effect was more pronounced when cells were exposed to PMB. Here, a sharp quenching followed by rapid dequenching was observed, indicating initial outer membrane permeabilisation followed by later inner membrane depolarisation. This is consistent with the two-step mode of action of PMB [46]. If the cells instead were pre-treated with PMBN, faster quenching was observed upon addition of DiSC_3_(5) (Fig. 5c). This was also accompanied by more extensive dequenching upon depolarisation induced by PMB addition.

**Figure 5:**
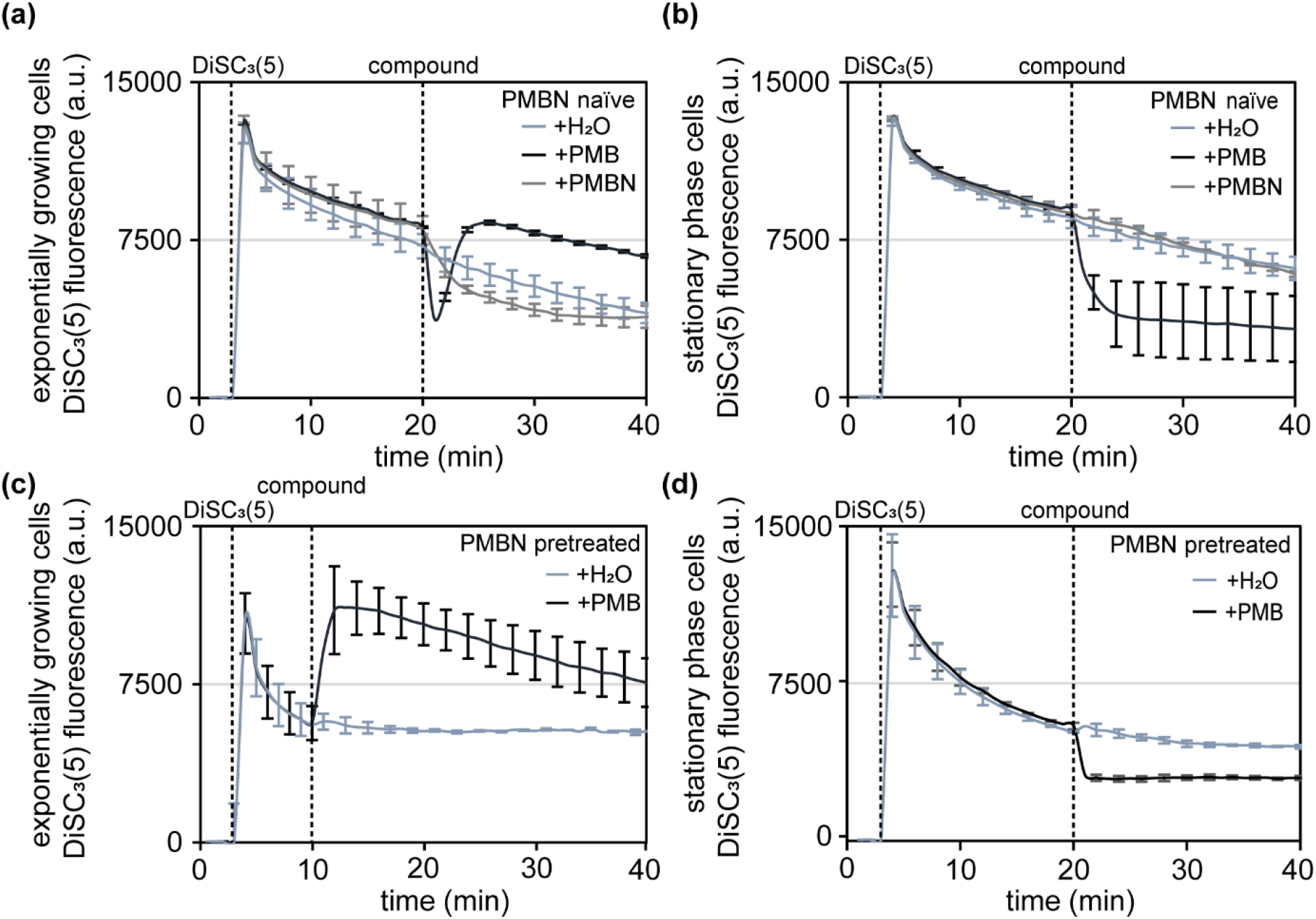
DiSC_3_(5)-based fluorometric measurement of membrane potential in *S. enterica.* PMBN naïve exponential (a) and stationary growth phase (b) *S. enterica* cells were exposed to either PMBN (30 μg/ml), PMB (20 μg/ml), or solvent (H2O). Dashed vertical lines indicate the addition of DiSC_3_(5) and compounds, respectively. Note the quenching of the DiSC_3_(5) fluorescence upon accumulation in cells, and rapid further reduction upon OM permeabilisation by PMB and PMBN followed by dequenching upon membrane depolarisation by PMB. (c-d) Measurements were repeated in cells pre-treated with PMBN. Graphs depict the means of six replicates and standard deviation from two independent experiments. Strain used *S. enterica* TH437 (wild type).

It is well-established that bacterial membrane potential can differ according to the growth phase or growth conditions. Whilst logarithmic growth phase cells feature well-energised membranes, entry into stationary growth phase is associated with nutrient limitations and other stresses that can lead to reduced membrane potential levels (also see movie S1) [47]. To establish that the fluorometric DiSC_3_(5) assay can also be applied for non-growing cultures, we repeated the experimental procedure with stationary phase cells obtained from an overnight culture. In the case of PMBN naïve cells, DiSC_3_(5) incorporation dynamics were similar for exponential and stationary phase cells (Fig. 5b), indicating that stationary phase *S. enterica* cells grown in rich medium maintain adequate membrane potential levels. Upon addition of PMB, a clear additional quenching step associated with outer membrane permeabilisation was observed. However, unlike in actively growing cells (Fig. 5a) this was not followed by dequenching associated with membrane depolarisation. Recently, Sabnis et al. demonstrated that inner membrane pore formation induced by Colistin (Polymyxin E) requires lipopolysaccharide (LPS), which is synthesised at the inner membrane prior to translocation to the outer membrane [7]. Very likely, the observed lack of depolarisation in non-growing cells is linked to this mechanism and caused by the absence of *de novo* LPS synthesis and, thus, inner membrane LPS. PMBN pre-treated stationary cells depicted a more rapid DiSC_3_(5) quenching behaviour compared to naïve cells, indicating that PMBN can permeabilise the outer membrane of stationary growth phase *S. enterica* to some degree (Fig. 5d). However, the inability of full length PMB to trigger additional dequenching in these cells, and the lack of a significant response when PMBN is added to naïve stationary growth phase cells (Fig. 5c) does suggest that PMBN might be less active against non-growing cells. Finally, to confirm that the established fluorometric assays are robust, we repeated the measurements for actively growing *E. coli* cells in a different laboratory setting and using different instrumentations. Indeed, very comparable results were obtained for *E. coli* (Fig. 6).

**Figure 6:**
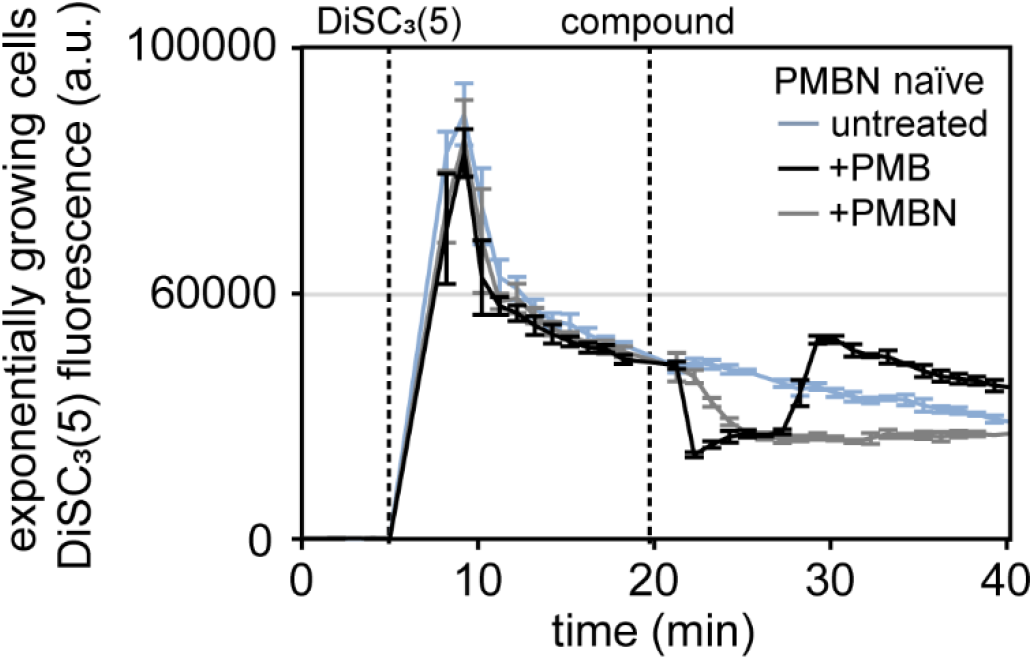
DiSC_3_(5)-based fluorometric measurement of membrane potential in *E. coli.* PMBN naïve exponential growth phase *E. coli* cells were exposed to either PMBN (30 μM) or PMB (7 μM). Dashed lines indicate addition of DiSC_3_(5) and compounds, respectively. Note the quenching of the DiSC_3_(5) fluorescence upon accumulation in cells, and further reduction upon OM permeabilisation triggered by PMB and PMBN, followed by dequenching upon membrane depolarisation triggered by PMB. The graph depicts mean and standard deviation from technical triplicates. Strain used *E. coli* MG1655 (wild type).

Whilst the influence of both outer membrane permeabilisation and cytoplasmic membrane depolarisation on DiSC_3_(5) persists in the fluorometric assay, pre-incubation with PMBN allows this confounding factor to be largely eliminated at least in case of actively growing cells. Hence, with careful controls, DiSC_3_(5) can be used to reliably monitor membrane potential in a microtiter plate format using a fluorometric approach.

## SUMMARY

Previously, we have published detailed methods and guidance on using carbocyanide-based voltage-sensitive dyes for the analysis of membrane potential in Gram-positive bacteria [23]. Due to the additional outer membrane and its impact on dyes and membrane-active compounds, translating those methods to Gram-negative bacteria is not necessarily trivial. In this report, we summarise our experiences using DiSC_3_(5) as a voltage-sensitive dye in Gram-negative species including *E. coli* and *S. enterica.* Whilst the use of DiSC_3_(5) as a reporter for membrane potential in Gram-negative bacteria is not novel, the information about best practices and factors that can compromise the measurements are not well documented, making the use of such dyes without prior knowledge and experience challenging. The methods presented here should be easy to implement using commonly available equipment such as regular widefield fluorescence microscopes and fluorescence plate readers. Furthermore, these assays should in principle be directly transferrable to flow cytometry measurements, although not verified within this study. We hope that the included details and discussions, and information regarding the effects and problems associated with outer membrane permeabilisation and the use of buffers rather than growth media, will provide a valuable starting point for those interested in analysing bacterial membrane potential in a physiological context, or as an assay to study antimicrobial mode of action.

## Abbreviations

DiSC_3_(5): 3,3’-Dipropylthiadicarbocyanine iodide
DiOC_2_(3): 3,3’-Diethyloxacarbocyanine Iodide
EDTA: Ethylenediaminetetraacetic acid
MIC: Minimal inhibitory concentration
ThT: Thioflavin T
PMB: Polymyxin B
PMBN: Polymyxin B nonapeptide
PBS: Phosphate-buffered saline; DMSO, Dimethyl sulfoxide

## Authors and contributors

PFP and HS designed and coordinated research; JAB, MH, PFP performed the experiments; JAB, PFP, and HS analysed data; JAB, PFP and HS wrote the paper; ME and HS acquired the funding. All authors commented on the manuscript.

## Conflicts of interest

The authors declare that there are no conflicts of interest.

## Funding information

H.S. was supported by UKRI (UK Research and Innovation) Biotechnology and Biological Sciences Research Council Grant (BB/S00257X/1) and J.A.B. by UKRI Medical Research Council Grant MR/N013840/1. This work was supported in part by a project that has received funding from the European Research Council (ERC) under the European Union’s Horizon 2020 research and innovation programme (grant agreement n° 864971) (to M.E.)

## Acknowledgements

We would like to acknowledge Maria Dakes Stavrakakis for preliminary work related to use of DiSC_3_(5) in stationary phase cells.

## Notes

### Competing Interest Statement

The authors have declared no competing interest.

