## Supplementary Information for "A guide for membrane potential measurements in Gram-negative bacteria using voltage-sensitive dyes"

Supplementary data

Authors: Jessica A. Buttress[1], Manuel Halte[2], J. Derk te Winkel[1], Marc Erhardt[2,3], Philipp F. Popp[2], and Henrik Strahl[1]

Affiliations:
[1]Centre for Bacterial Cell Biology, Biosciences Institute, Faculty of Medical Sciences, Newcastle University, Newcastle upon Tyne, UK
[2]Institute for Biology - Bacterial Physiology, Humboldt-Universität zu Berlin, Berlin, Germany
[3]Max Planck Unit for the Science of Pathogens, Berlin, Germany.

Content:

Supplementary Figure 1
Supplementary Figure 2

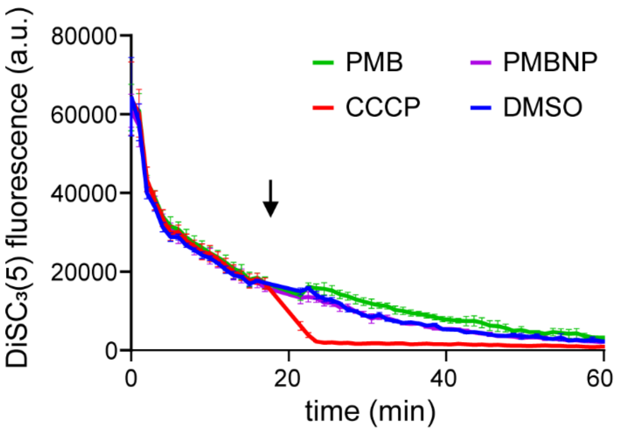


**Supplementary Figure 1: Interactions between DiSC$_3$(5) and antimicrobial compounds.** Fluorescence intensity of 1 µM DiSC$_3$(5) in PBS supplemented with 0.5 mg/ml BSA upon addition of the pore forming antibiotic PMB and the outer membrane permeabilising agent PMBN. The time point of compound addition is indicated by an arrow. Addition of the protonophore CCCP is shown as an example of strong dye interference. The graph depicts the mean and standard deviation of technical triplicate measurements.

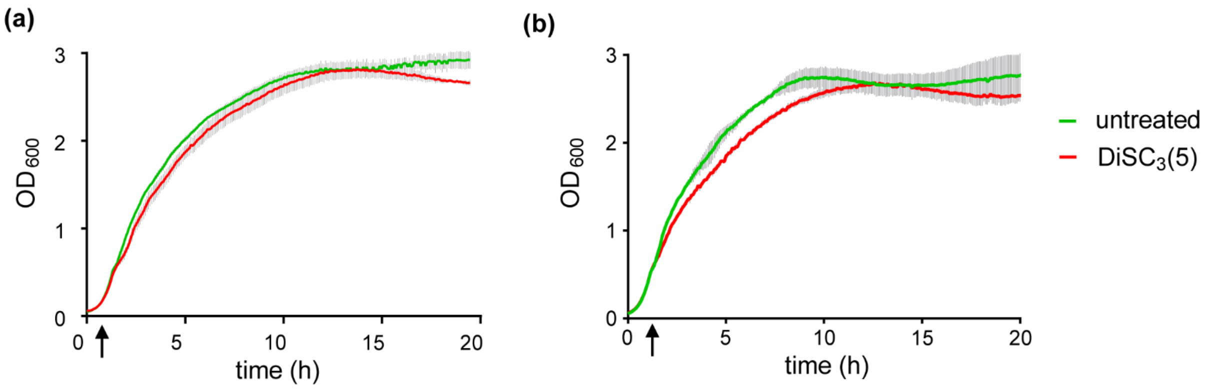


**Supplementary Figure 2: $DiSC_3(5)$ does not inhibit *E. coli* growth.**

Growth curve of *E. coli* cells grown to $OD_{600}$ 0.5, followed by addition of 1 µM $DiSC_3(5)$ in the absence (a) and presence (b) of PMBN (30 µM). $DiSC_3(5)$ addition is indicated with by an arrow. The graph depicts the mean and standard deviation of technical triplicate measurements. Strain used: *E. coli* MG1655 (wild type).
